# Overlooked polyploidies in lycophytes generalize their roles during the evolution of vascular plants

**DOI:** 10.1101/217463

**Authors:** Jinpeng Wang, Jigao Yu, Pengchuan Sun, Chao Li, Xiaoming Song, Tianyu Lei, Yuxian Li, Jiaqing Yuan, Sangrong Sun, Hongling Ding, Xueqian Duan, Shaoqi Shen, Yanshuang Shen, Jing Li, Fanbo Meng, Yangqin Xie, Jianyu Wang, Yue Hou, Jin Zhang, Xianchun Zhang, Xiyin Wang

## Abstract

Seed plants and lycophytes constitute the extant vascular plants. As a model lycophyte, *Selaginalla moellendroffii* was deciphered its genome, previously proposed to have avoided polyploidies, as key events contributing to the origination and fast expansion of seed plants. Here, using a gold-standard streamline recently proposed to deconvolute complex genomes, we reanalyzed the *S. moellendroffii* genome. To our surprise, we found clear evidence of multiple paleo-polyploidies, with one being recent (~ 13-15 millions of years ago or Mya), another one occurring about ~125-142 Mya, during the evolution of lycophytes, and at least 2 or 3 events being more ancient. Besides, comparison of reconstructed ancestral genomes of lycophytes and angiosperms shows that lycophytes were likely much more affected by paleo-polyploidies than seed plants. The present analysis here provides clear and solid evidence that polyploidies have contributed the successful establishment of all vascular plants on earth.

## Introduction

Extant vascular plants can be divided in two types, the duphyllophytes (ferns and seed plants) and the lycophytes, which had diverged as early as 410 million years ago (Banks et al. 2011). As a model plant lycophyte, *Selaginalla moellendorffii*, featuring a dominant and complex sporophyte generation and vascular tissues with lignified cell types, was deciphered its genome sequence, ~ 212.6 Mbp containing ~ 22,285 genes(Banks et al. 2011).

Recursive polyploidies, or whole-genome duplications, have been proposed to be a key evolutionary drive force of seed plants, responsible for their divergence and fast expansion on earth(Paterson et al. 2004; Jiao et al. 2011). Considerably different from seed plants, it was reported that the *S. moellendorffii* genome lacks evidence of any paleo-polyploidies(Banks et al. 2011).

Plant genomes can be quite complex, at least partially due to recursive polyploidies and accumulation of repetitive sequences. It is often difficult to perform a comprehensive analysis to understand their genome structure and evolution.

There occurred quite several times that certain ancestral polyploidies eluded from genome analysis, resulting in problematic interpretation of the structure, evolution, and/or functional innovation of whole genomes and key gene families(Paterson et al. 2005; Wang et al. 2005; Li et al. 2015; Wang et al. 2016b). In sake of much time, energy, and money invested in a genome project, this can be a nonelected pity.

Recently, we proposed a gold standard streamline to deconvolute complex genomes, esp. of plants, and using it reanalyzed the cucubit genomes and revealed an overlooked paleo-tetraploidy, occurring ~100 Mya, in the common ancestor of cucurbiticeae plants(Wang et al. 2017a).

Here, we used the streamline to perform a comprehensive analysis of the *S. moellendorffi* genome and other plant genomes. Notably, we came to the findings of multiple polyploidies during the evolution of the lycophytes. Further comparative genomics analysis suggested general roles of polyploidies during the early origination and divergence of vascular plants.

## Results

Using a gene-colineartiy based approach implemented in ColinearScan, with maximal gap of 50 genes between neighboring colinear genes, we inferred 302 syntenic blocks in *S. moellendorffii* genome **(Supplemental Tables S1 and S2)**.

These blocks involved 2, 632 colinear gene pairs, spanning genomic regions of an accumulated 154.13 Mbp (72.50% of 212.6Mbp) **(Fig. 1A)**. They covered 86.33% (19, 239/22, 285) of all genes, and 87.55% (19, 239/21, 975) of assembled genome sequences, related to all 361 scaffolds, which at least 53.19% (192/361) were covered by syntenic blocks with 5 or more colinear genes.

**Figure 1A.**
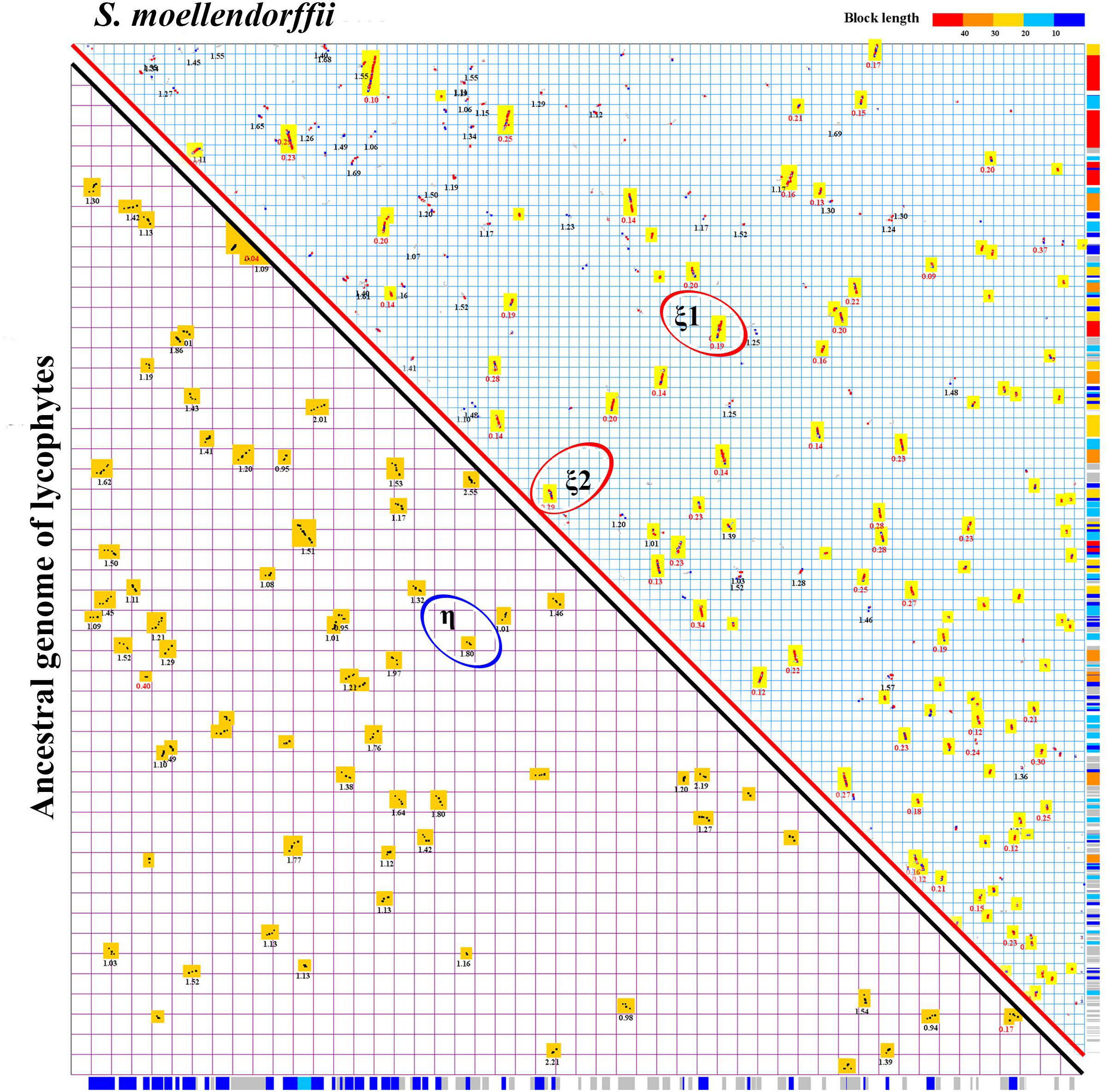
Homologous gene dotplot of *S. moellendroffii*. The upper-right diagonal shows dotplots between scaffolds of extant genome. Both X and Y axes represented by sequentially linked scaffolds. A duplicated gene pair inferred by ColinearScan between X and Y results in a dotplot in the figure. Neighboring dotplots resulted in blocks showing duplicated segments in the genome. Dotplots of the best hit genes are in red, the secondary ones in blue, and others in grey. The median Ks value of gene pairs in a block is displayed, and colored in red when Ks < 0.5, in black when Ks > 1 and Ks < 1.7. The blocks in the recent tetraploidization, highlighted in yellow boxes, are mapped to the Y-axis, with longest blocks colored in a neighboring region displayed with a color scheme showed along Y-axis in the right The lower-left shows dotplots in the reconstructed ancestral genome before the recent tetraploidy. Dotplots are produced between genes in the ancestral genome. Blocks are highlighted and Ks values of blocks are displayed. The blocks are mapped to the X, with the same color scheme defined above to show longest blocks in a neighboring region. Examples of duplicated blocks produced by η and ξ are circled out, to display gene colinearity in subfigure **1B**.

The syntenic blocks were mapping onto the chromosomes, and produced coverage as deep as 10. Around 19.24% of genome regions were covered in a depth of 4 or more, and 17.55%, 25.36%, 27.06%, 10.79% with 3, 2, 1, 0 **( Supplemental Fig. S1 and Supplemental Table S3)**. Articulated correspondence of colinear genes between multiple duplicated regions could often be found, showing likely recursive gene duplications **(Fig. 1B)**.

**Figure 1B.**
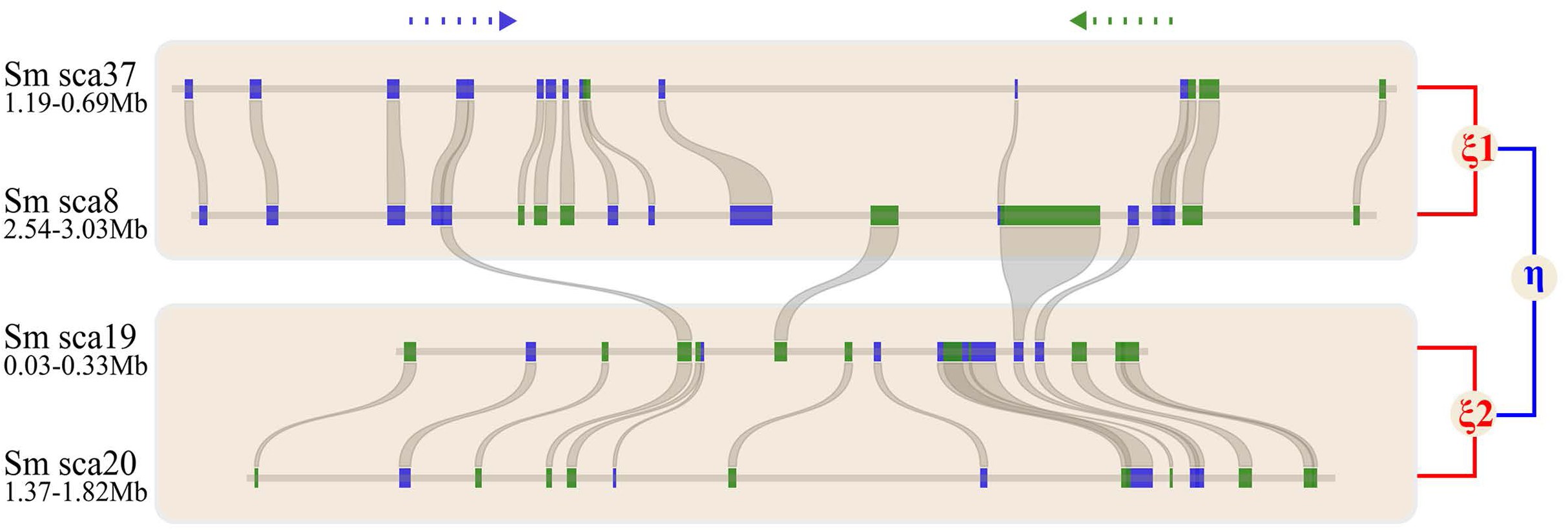
Alignment of genomic regions with colinear genes. The involved four regions are from *S. moellendroffii* scaffold20, scaffold19, scaffold8, and scaffold37, inferred to have been produced by two recursive polyploidies, η and ξ. Genes are shown with rectangles, and different colors show transcription directions. Colinear genes were linked with curvy lines.

Diverged genic sequence divergence of colinear genes clearly showed the occurrence of two polyploidies. We characterized the synonymous nucleotide substitutions at all or only four-degenerate synonymous substitution sites or only four-degenerate transversions on the third codon sites, namely Ks and 4Dtv, and the distribution of them both displayed a clear bi-modal distribution **(Fig. 2 and Supplemental Fig. S2)**. As to Ks, the two peaks were located at 0.12 and 1.2 respectively. Based on Ks distribution, we divided the colinear genes into two groups **(Fig. 1A)**. Supposing a 6.0-7.0 × 10^−9^ synonymous substitutions per site per year, borrowed from angiosperms(Moniz and Drouin 1996; Paterson et al. 2004), two peaks were re-posited at 0.12 and 1.15, respectively, and two large-scale genomic duplication events can be inferred to have occurred ~13-15 millions of years ago (mya, named ξ or *Xi*), and ~125 - 142 (mya, named η or *Eta*). Using separate distributions rather than the merged distribution could eliminate interactive effect during statistical analysis. The younger group covered 58.41%(13, 017/22, 285 genes) of genome, and was related to all chromosomes and 70.31% (135/192) of the scaffold, showing that ξ was a polyploidization. By merging the colinear genes of younger group, we reconstructed the ancestral genome content before the event. Then, we inferred 120 syntenic blocks in the reconstructed ancestral genomes, which involved 936 colinear genes and the spanning regions covered 73.17% (112.78Mbp) of the ancestral genome, showing that likely η had also been a polyploidization.

**Figure 2.**
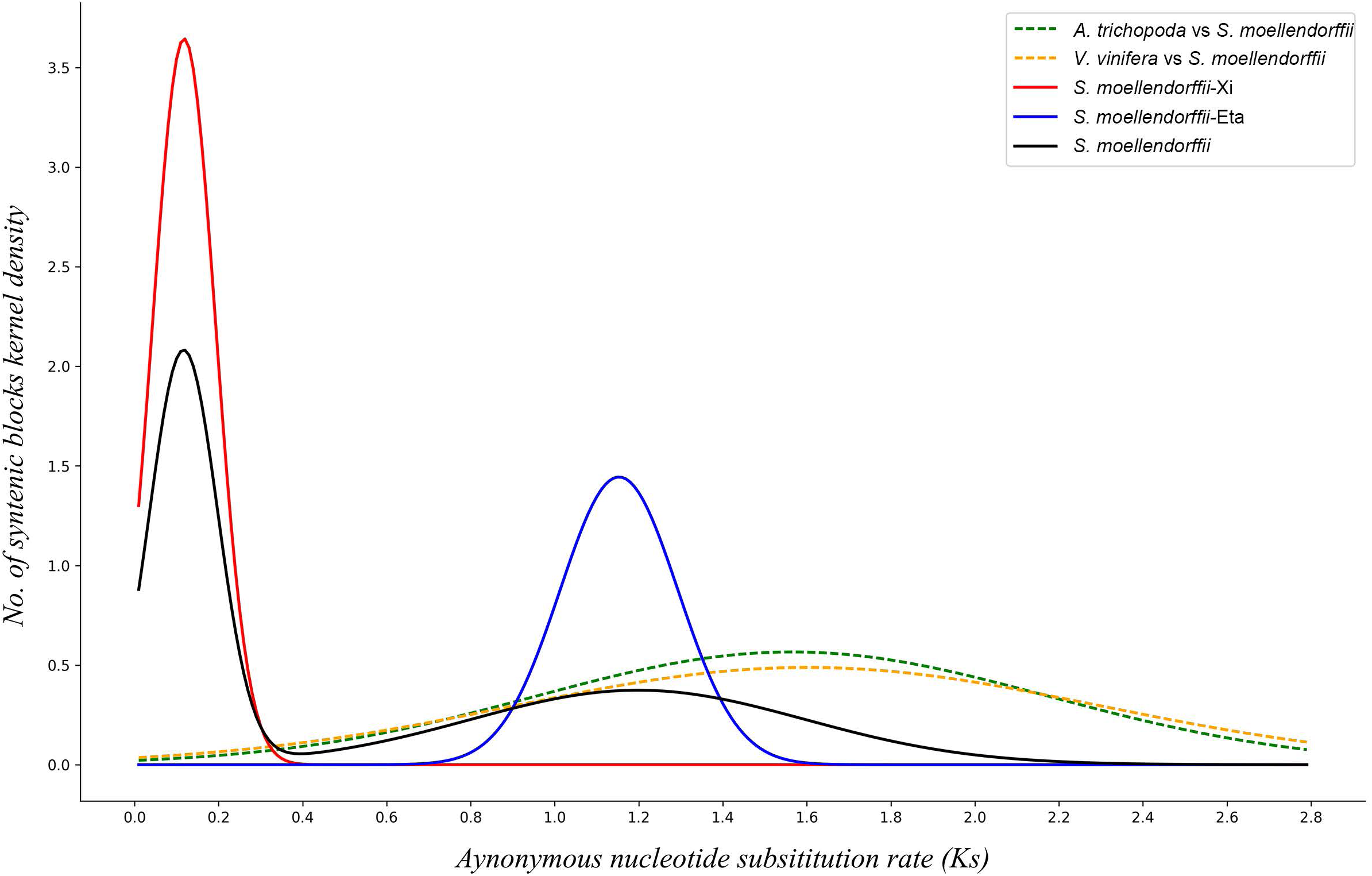
Distribution of Ks between colinear genes.

Furthermore, we checked whether the ancient event was shared with seed plants. By inferring colinear blocks and characterizing Ks between colinear genes between *S. moellendorffii* and two model angiosperms, *Amborella trichopoda* and *Vitis vinifera* (grape), respectively, we found that Ks peaks of putative orthologous genes at 1.56 and 1.6 **(Fig. 2)**. This suggested that above two events inferred here both occurred in the lycophyte lineage after their split from seed plants **(Fig. 3)**.

**Figure 3.**
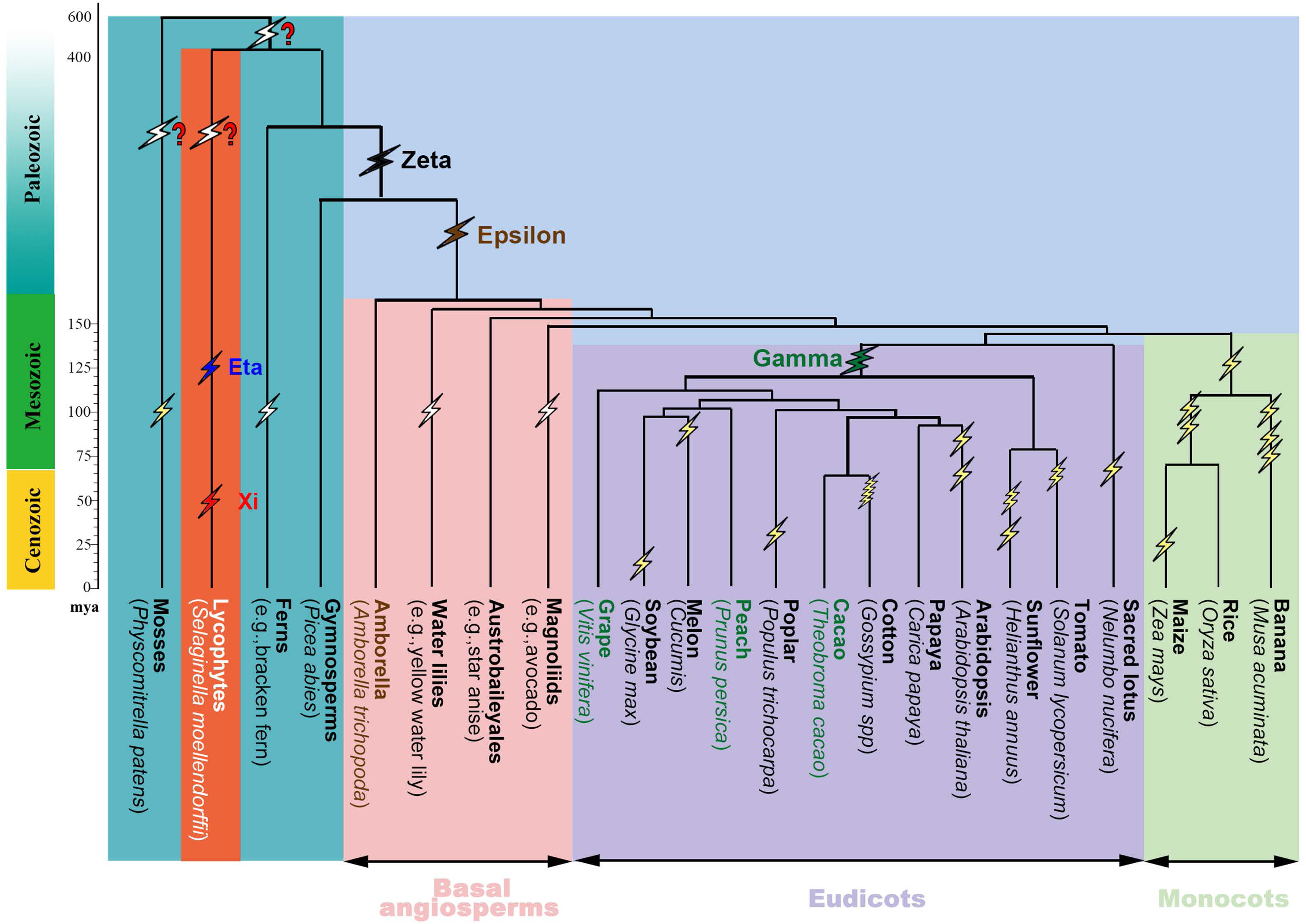
Plant phylogeny and inferred ancient polyploidies. Flashlight marks are used to show polyploidy events, with those with two-turn zigzag to show a tetraploidy, the three-turn turn ones to show a hexaploidy, and five-turn ones to show decaploidy, and those with a question mark to show non-determined ploidy level. The events, η and ξ, inferred in the present work are shown blue and red, respectively. More ancient events that were inferred to have likely occurred were also shown with question marks.

An appreciable fraction of *S. moellendorffii* genomic regions were covered in depth > 4 by colinear blocks, suggesting the likelihood of more ancient polyploidies. Therefore, we performed a deeper search of gene synteny between *S. moellendorffii* and two referenced angiosperm genomes. The Amborella genome, assembled into scaffolds, has a simple structure, avoiding a genome doubling after the split with other angiosperm and providing evidence of a more ancient polyploidy. The grape genome, being well assembles into pseudochromosomes, we involved it and reconstructed the ancestral genome before a major-eudicot-common hexaploidy.

We inferred syntenic blocks between *S. moellendorffii* and each of the referenced genomes, and mapped the blocks onto each genome **(Supplemental Fig. S3 and Fig. S4)**. As to the *S. moellendorffii –*Amborella homology, 14.18% and 9.74% of *S. moellendorffii*, and Amborella genomes, respectively, were covered by colinear blocks to a depth of 4 or more **(Supplemental Table S4)**. As to *S. moellendorffii –* grape homology, 9.92% and 6.58% of the *S. moellendorffii* and grape genomes were covered by colinear blocks to a depth of 4 or more **(Supplemental Table S5)**. These finding suggested more polyploidies during the evolution of vascular plants. Then, we explored the homologous region coverage depth in reconstructed the pre-ξ ancestral genome of lycophytes (ALG, 11509 genes), and that of angiosperms (AAG, 1686 genes) inferred based on the grape genome. The ALG involved genes from 34 largest *S. moellendorffii* scaffolds. At the gap size 50 genes when running the ColinearScan, the significant colinear blocks with at least 4 colinear genes resulted in homologous coverage depth as deep as 7 and 18 in the AAG and ALG, respectively **(Fig. 4)**. As to coverage depth, 78.3% and 7.5% of the AAG and ALG were covered to depth 4 and more **(Supplemental Table S6)**. These findings implied more than 2 or more ancient polyploidies beyond the ancient polyploidy in lycophytes that we found above, and the polyploidy revealed with Amborella genome in the seed plant lineage(Project et al. 2013). Moreover, a more coverage onto the AAG genome means that lycophytes might have experienced many more polyloidies than seed plants have. To be careful, when searching colinear blocks, we tried different maximal gap sizes between neighboring colinear genes in each lineage **(Supplemental Tables S6B-K)**. Though homologous gene coverage varied, we came to similar conclusion **(Fig. 3)**. Further determination of numbers and dates of these potential events has to be left for future research by involving more taxa.

**Figure 4.**
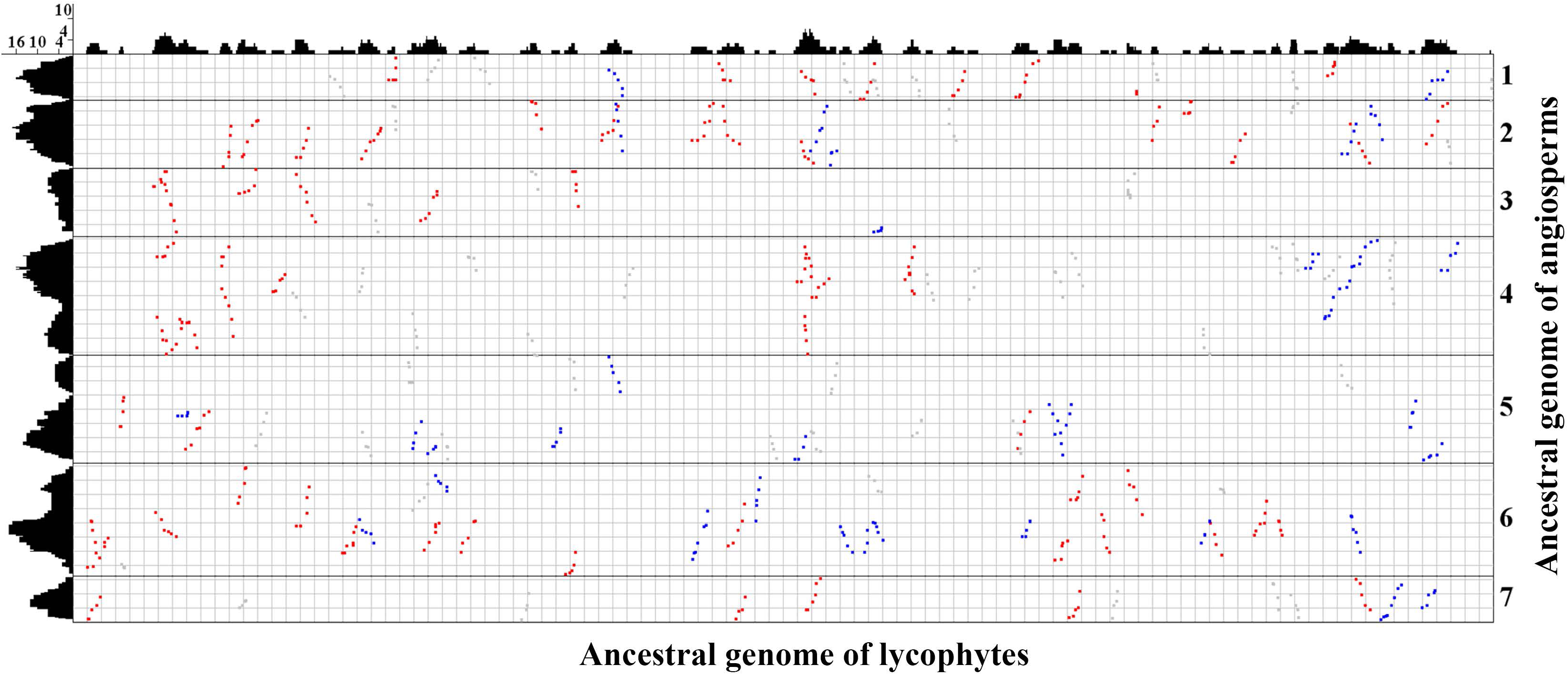
Inferred colinear genes between ancestral lycophyte and angiosperm genomes, i.e., ALG and AAG. Inferred ancestral regions represented by genes were arranged along X and Y axe, respectively. Statistically significant colinear blocks are displayed, and mapped onto each axis to produce homologous coverage depth. Blocks with median Ks <1.8 colored by red; Blocks Ks >= 1.8 colored by blue; others colored by grey.

## Discussion

Based on phylogenetic tree construction, polyploidies was inferred to have contributed to the early divergence of seed plants(Jiao et al. 2011). By dissecting features of Ks distribution with transcriptome data, polyploidy was inferred to have affected a basal land plant, horsetail(Vanneste et al. 2015). A large-scale of cytogenetic and phylogenetic analysis indicated that ferns might have been more frequently affected by polyploidies(Wood et al. 2009). Together with many other publications proposing ancient polyploidies(Soltis et al. 2009; Peer 2011; D’Hont et al. 2012; Chalhoub et al. 2014), these works each shed light on the far past of plant evolution. However, though the Seleginalla genome has been ready for six years, a genome-scale evidence of ancient polyploidies in the lycophytes has still been elusive. Here, using a gold-standard streamline deciphering complex genomes, our findings provide clear and solid evidence that two polyploidies have contributed to evolution of the lycophytes after their split with seed plants, and several older ones are common to all vascular plants.

The availability of whole-genome sequences provides evidence of ancient polyploidies that one can *see* other than *infer*. However, as mentioned previously, there have been quite some failures in deconvoluting these large-scale events(Paterson et al. 2005; Wang et al. 2005; Li et al. 2015; Wang et al. 2016b). The initial analysis of Selagillana genome overlooked these large-scale evolutionary events. The reasons resided in the complicated nature of plant genomes, drawbacks in adopted methodology, and/or inexperienced bioinformatics teams especially in some genome-sequencing companies, as discussed previously (Jinpeng Wang 2017). Compared to the whole-genome approach, inferences based on statistical or computational approaches dissecting complex data such as Ks distribution or phylogenetic tree topology, provide much weaker evidence. These facts highlight the importance of the present whole-genome level reanalysis of the Selaginalla genome using the gold-standard streamline to understand complex genomes.

Further research will help understand how these polyploidies have contributed to the biological and genetic innovations during the evolution of vascular plants, especially, lycophytes.

## Methods

The involved genome sequences and annotations were from public database **(Supplemental Table S7)**. Colinear genes were inferred by using ColinearScan(Wang et al. 2006), a well statistically supported algorithm and software. Maximal gap length between genes in colinearity along a chromosome/scaffold/contig sequence were set to be 50 genes apart, which have been adopted in many previous publications(Wang et al. 2015; Wang et al. 2016a; Wang et al. 2016b; Liu et al. 2017; Wang et al. 2017b). Homologous gene dotplots within a genome or between different genomes were produced by using MCSCAN toolkits(Wang et al. 2012), which the corresponding author contributed to direct development. The gold-standard streamline to decipher complex genomes was well followed as described previously (Jinpeng Wang 2017). Evolutionary divergence between homologous genes was estimated as previously(Wang et al. 2015; Wang et al. 2017b).

Construction of ancestral genomes was also based on our previously detailed methods (Jinpeng Wang 2017). For clarification, here, we reconstructed the ancestral lycophyte genome before the polyploidy ξ (ALG), and the angiosperm ancestral genome (AAG). The ALG genome was reconstructed by using one copy of the colinear genes produced by the recent polyploidy, in that the colinear genes probably preserved their ancestral gene location **(Supplemental Fig. S5A)**. In a similar approach, the AAG genome was reconstructed by inferring ancestral genes using grape colinear genes preduced by the major-eudicot-common hexaploidy (previously named ϒ) **(Supplemental Fig. S5B)**. We did not infer AAG by using grape-Amborella or grape-rice comparison due to incomplete assembly of Amborella genome or severe genomic fractionation after multiple polyploidies in monocot lineage(Tang et al. 2010).

## Acknowledgements

We appreciate financial supported from the Ministry of Science and Technology of the People’s Republic of China (2016YFD0101001), China National Science Foundation (31371282 to X.W. and 3151333 to J.W.), Natural Science Foundation of Hebei Province (C2015209069 to J.W.). and Tangshan Key Laboratory Project to X.W.; Hebei New Century 100 Creative Talents Project, Hebei 100 Talented Scholars project, and Tangshan Key Laboratory Project to X.W. We thank the helpful discussion with researchers at the iGeno co Ltd, China.

## Author contributions

X.W. conceived and led the research. J.P.W. implemented and coordinated the analysis. J.Y., P.S., C.L., X.S., Y.L., J.Y., S.Sun, H.D., X.D., S.Shen, Y.S., J.L., F.M., Y.X., J.Y.W., Y.H., J.Z. performed the analysis. J.Y. and T.L. contributed analyzing tools. X.W. and wrote the paper. X.Z. contributed to plant phylogeny.

